# High-Resolution Single-Molecule FRET via DNA eXchange (FRET X)

**DOI:** 10.1101/2020.10.15.340885

**Authors:** Mike Filius, Sung Hyun Kim, Ivo Severins, Chirlmin Joo

## Abstract

Single-molecule FRET is a versatile tool to study nucleic acids and proteins at the nanometer scale. However, currently, only a couple of FRET pairs can be reliably measured on a single object. The limited number of available FRET pair fluorophores and complicated data analysis makes it challenging to apply single-molecule FRET for structural analysis of biomolecules. Currently, only a couple of FRET pairs can be reliably measured on a single object. Here we present an approach that allows for the determination of multiple distances between FRET pairs in a single object. We use programmable, transient binding between short DNA strands to resolve the FRET efficiency of multiple fluorophore pairs. By allowing only a single FRET pair to be formed at a time, we can determine the FRET efficiency and pair distance with sub-nanometer resolution. We determine the distance between other pairs by sequentially exchanging DNA strands. We name this multiplexing approach FRET X for FRET via DNA eXchange. We envision that our FRET X technology will be a tool for the high-resolution structural analysis of biomolecules and other nano-structures.

## INTRODUCTION

X-ray crystallography, nuclear magnetic resonance (NMR) and cryo-electron microscopy are the golden standard for determining the structure of biomolecules.^1,2^ However, minute and rapid conformational changes often cannot be observed with these techniques, since the required sample preparation may stabilize a certain conformation of a molecule.^3^ In addition, none of these techniques are capable of detecting rare species in the cell, for which single-molecule sensitivity is required. Single-molecule FRET can be used to determine the structure of molecules with sub-nanometer resolution. However, the use of single-molecule FRET for the analysis of complex molecular structures (e.g. protein tertiary structures) has been limited since it requires resolving the FRET efficiency of multiple dye pairs.^4,5^ Currently, single-molecule FRET analysis allows us to deal with only one or two FRET pairs in a single measurement.^6,7^ Therefore, structural analysis using single-molecule FRET requires the preparation of a protein library consisting of many different combinations of dye locations, rigorous modeling and simulations following the data acquisition. ^8–11^

Single-molecule multiplexing has been demonstrated with photoswitchable fluorophores. In this approach, a molecule of interest is labeled with a single donor and two or more identical acceptor fluorophores. By using photoswitchable acceptor fluorophores, only one of the acceptors is active at a given time.^12^ This method, called photoswitchable FRET, allows for the detection of multiple FRET pairs in a single nanoscale object and determination of structures and interactions between biomolecules, from proteins to DNA. However, the stochastic nature of the photoswitching and the limited number of orthogonal attachment chemistry strategies for dye labeling are the main obstacles for the wide adaptation of the method. An alternative way of switching between on and off states of fluorescent probes is by using fluorophores that bind a target only for short period of time, as with point accumulation in nanoscale topography (PAINT).^13–15^ For example, fluorophores are attached to short DNA oligos that bind the complementary target strands for several hundreds of milliseconds. This transient binding is central to the super-resolution technique, DNA-based point accumulation for imaging in nanoscale topography (DNA-PAINT).^16–19^

Here we propose a new single-molecule structural analysis tool that can resolve the FRET efficiency of multiple pairs in a single target molecule. By using programmable, transient binding between short DNA strands, a single FRET pair is formed at any given time allowing for distance determination between the momentarily activated fluorophore pair. By repeating the imaging cycle, we can resolve multiple points of interest (POI) in a single nanoscale object. We demonstrate the proof of concept of sub-nanometer resolution single-molecule structural analysis on DNA structures.

## RESULTS

To demonstrate the concept of FRET via DNA imager strands, we first tested single molecule FRET measurements with a transiently binding dye-labeled DNA imager strand. We designed an assay where an acceptor (Cy5)-labeled single-stranded (ss) DNA strand is immobilized on a quartz slide through biotin-streptavidin conjugation (**Figure 1A**). The measurements yielded a distinct FRET signal upon binding between a donor-labeled imager strand and the immobilized target strand (**Figure 1B**). The FRET signals were recorded using total-internal-reflection microscopy. The imager strand sequence was chosen such that the binding events between the two DNA strands would have a short dwell-time to allow for frequent replenishment of the imager strand (**Figure 1C**). This allows for the same POI to be probed multiple times. At the same time, the dwell-time of the binding events between the imager and immobilized strands was chosen to be several hundred milliseconds or longer for precise determination of the FRET efficiency.

**Figure 1:**
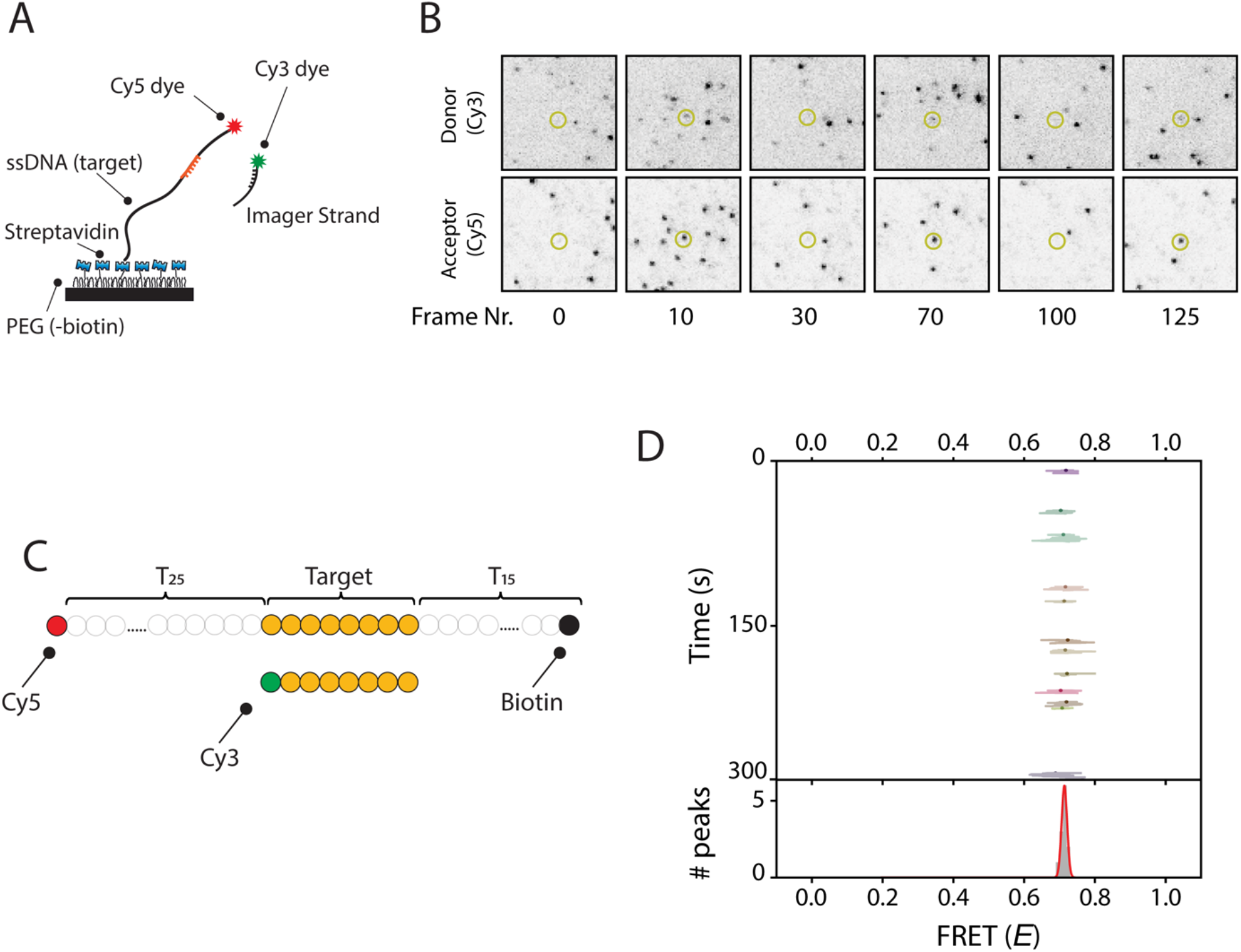
Repetitive binding of short DNA imager strand allows for high-detection precision for Single Molecule FRET. A)Schematic representation of the single-molecule FRET assay. An acceptor (Cy5, red star) labeled single-stranded target DNA construct is immobilized on a PEGylated surface through biotin-streptavidin conjugation. Binding of the donor (Cy3, green star) labeled imager strand results in short FRET events and is observed using total internal reflection microscopy. B)A series of CCD snapshots obtained from a single-molecule movie with 100 ms exposure time. The top row represents the donor channel, and the bottom row represents the acceptor channel. Each dot represents a single molecule. Dynamic binding of the imager strands can be observed over time (highlighted molecule). C)Schematic representation of the ssDNA constructs. Upon binding of the imager strand, the donor fluorophore is separated from the acceptor by a 25-nt thymine linker. D)Single-molecule FRET kymograph from a time trace from one single molecule (highlighted molecule from Fig. 1B). The kymograph shows the FRET efficiency for each data point in a binding event (lines) and the mean FRET efficiency from all data points per binding event (dots) as a function of time. All mean FRET efficiencies are plotted in a histogram and fitted with a Gaussian function (bottom panel).

To visualize the FRET efficiency of each dye pair appearing in a field of view, we built a FRET kymograph (**Figure 1D and Supplementary Figure 1A**). The kymograph shows the FRET efficiency per data point (**Figure 1D, lines**) and the mean FRET efficiency from all data points per binding event (**Figure 1D, dots**). A FRET histogram built from the mean values is fitted with a single Gaussian distribution (**Figure 1D, bottom**) (*E* = 0.72 with a standard deviation of 0.01). The ensemble kymograph built from all 363 molecules for this construct shows a similar mean FRET of 0.71 ± 0.02 (**Supplementary Figure 1B**).

Structural analysis of complex biomolecules using single molecule FRET requires the detection of multiple FRET pairs in a single object. To avoid the crosstalk between different FRET pairs, each POI should be measured for a short amount of time using short DNA imager strands, thereby separating the FRET binding events of each pair in time. Therefore, we designed donor-labeled imager strands that can interact with a single POI only for 2-3 seconds (**Supplementary Figure 2A and B**), which is long enough to determine the FRET efficiency. We designed a ssDNA construct with two target sequences. The binding of a donor-labeled imager strand to the ssDNA construct yielded either a high or a medium FRET signal (**Figure 2A**). For a target construct that consisted of two POIs that were spaced by a 5-nt linker (**Figure 2B**), two FRET peaks were observed (**Figure 2D**), reporting on the location of each POI. However, when the two POIs were placed with no linker sequence in between (**Figure 2C**), the FRET histogram became unresolvable (**Figure 2E**). We conclude that it is not feasible to determine the pair distances of several POIs with high spatial resolution using a single imager strand.

**Figure 2:**
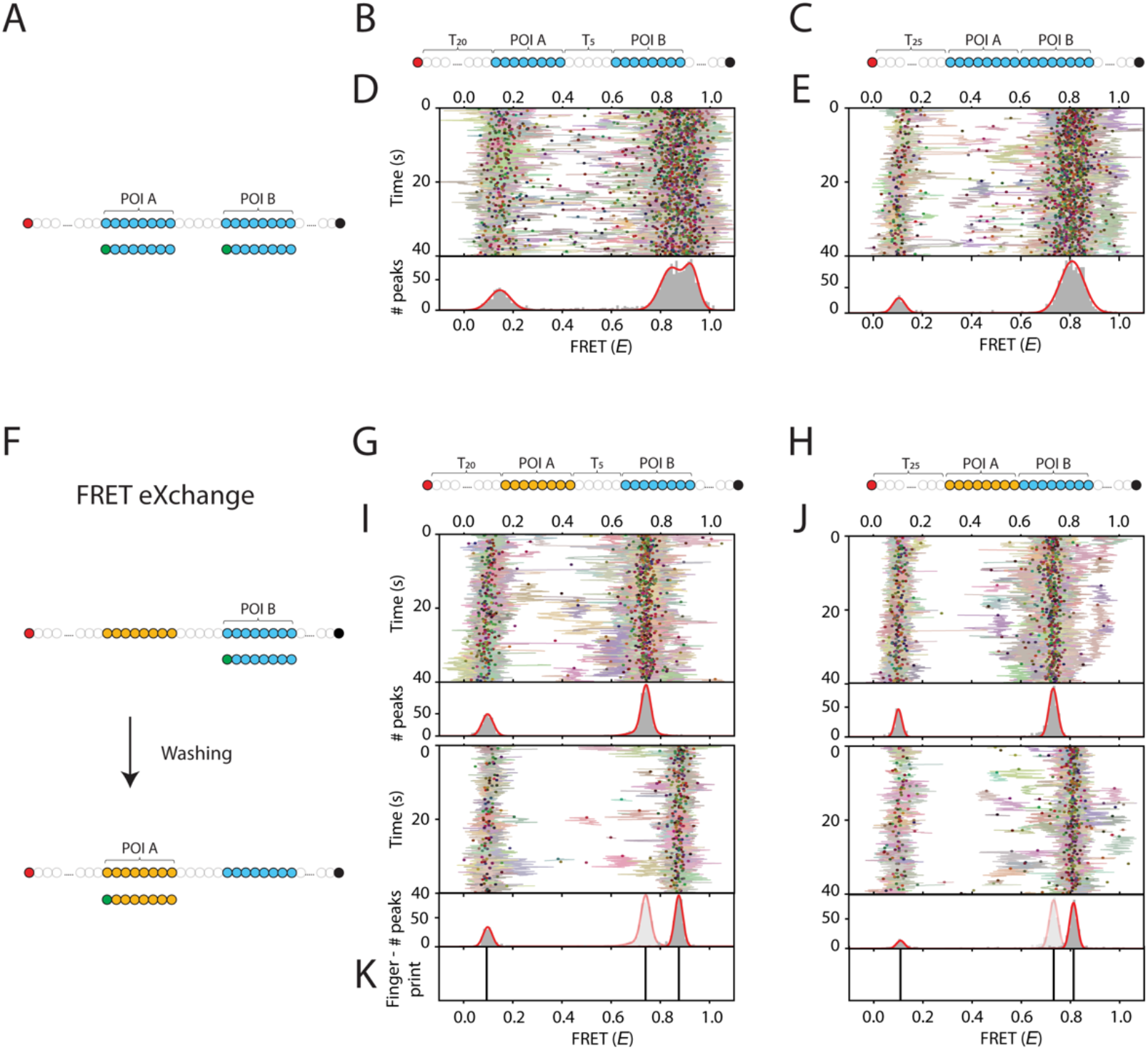
FRET by eXchange of unique imager strands allows for high spatial resolution of multiple POIs in a single nanoscale object. A) Schematic representation of the single-molecule experiments with two target sequences. A single imager strand is used that can bind to both of the POIs in a target molecule. An acceptor (Cy5, red circle) labeled ssDNA construct contains two POIs. Binding of the donor (Cy3, green circle) labeled imager strand results in either high FRET (when binding to POI A) or mid FRET (when binding to POI B). B and C) Schematic representation of the target constructs in which two POIs are separated by a 5 Thymine linker (Fig. 2B) or in which the two POIs are directly connected to each other (Fig. 2C). Single-molecule kymograph of the ssDNA target construct from Fig. 2B. Top panel shows the binding events obtained for all molecules in a single field of view. Bottom panel shows a FRET histogram consisting of a donor-only peak and two additional FRET peaks reporting on the location of each POI with respect to the acceptor fluorophore. The single-molecule kymograph of the ssDNA target construct from Fig. 2C. Using the same imager strand for both POI does not allow for the detection of the position of both POIs when they are in close proximity. The FRET histogram shows a broad peak at 0.81. Schematic workflow of FRET by eXchange of imager strand (or FRET X). A ssDNA target constructs consists of two POIs with unique DNA binding sequences, allowing us to measure the POIs one at a time. In a first round of detection, the imager strand for POI B (blue circles) is added and imaged for 5 minutes. Then the microfluidic chamber is washed and an imager strand for POI A (orange circles) is added. G and H) Schematic representation of the FRET X target constructs, in which two unique POI sequences blue circles (POI B) or orange circles (POI A) are separated by a 5 nt thymine linker (Fig. 2G) or in which the two POIs are directly adjacent (Fig. 2H). I and J) Single molecule kymographs for the FRET X for constructs in Fig. 2G and 2H. FRET X imaging allows for the determination of each POI in a separated round. For a construct in which the POIs were separated by a 5 nt thymine linker we observed a distinct FRET peak at 0.73 (Fig. 2I, top panel) and 0.88 (Fig. 2I, bottom panel) for POI A and B, respectively. FRET X allows for the accurate detection of POIs even when they are in closer proximity. We observed distinct FRET peaks of 0.73 (Fig. 2J, top panel) and 0.81 (Fig. 2J, bottom panel) for POI A and B, respectively. K) The Gaussian fits of individual histograms for each POI obtained using the FRET X approach allows for the determination of the center of a peak with 1 %p (or ΔE~0.01) precision. The centres of the peaks are plotted in a separate panel, which we name the FRET fingerprint of a nanoscale object.

To achieve higher spatial resolution, we sought to detect the different POIs independently so that the overlapping FRET peaks can be obtained separately and fitted more precisely. As illustrated in **Figure 2F**, each POI was measured using a unique short DNA imager strand. After the binding events for the first POI were recorded for several minutes, the imager strand was exchanged by washing the microfluidic chamber and injecting a unique DNA imager strand for the second POI (**Figure 2F**). This process can be repeated for any number of POIs. We name this “FRET X” for FRET via DNA eXchange.

To demonstrate the concept of FRET X, we designed ssDNA constructs with two POIs, each containing a unique target sequence. The POIs were separated by a 5-nt thymine (**Figure 2G**) linker or were in closer proximity with no linker in between (**Figure 2H**). In the first round of FRET X detection we determined the FRET peak to be at 0.74 for POI B located 35 nucleotides away from the acceptor (**Figure 2I top panel**). Next, the microfluidic chamber was washed and the imager strand complementary to the POI A was injected. In the second round of FRET X imaging, we observed a single FRET peak at 0.88 reporting on the second POI that was separated from POI B by a 5-nt thymine linker (**Figure 2I, bottom panel**). As shown in **Figure 2H**, FRET X allows for the accurate detection of the FRET efficiencies of both POIs, even when they not separated by a linker and this are in closer proximity. These FRET peaks could not be distinguished using conventional FRET, as shown in **Figure 2E**. In the first round of FRET X we observed a FRET peak at 0.73 for POI B (**Figure 2J, top panel**) and in the second round we observed a FRET peak at 0.81 for POI A (**Figure 2J, bottom panel**). Each histogram showed a wide distribution of ~ 10%p (the standard deviation) of the peak. However, the Gaussian fit can be used to resolve the center of a peak with high accuracy of ~1%p (standard error of mean) and is dependent on the number of binding events (**Supplementary Figure 3**). The resolved FRET values for each POI can be plotted as the FRET fingerprint of the measured object (**Figure 2K**).

To further investigate the achievable resolution of FRET X, we designed a series of imager strands in which the position of the donor fluorophore is altered by only a single base among the different imager strands (**Figure 3A**). The FRET X cycle was then repeated for all nine imager strands. The center of a peak of each histogram was determined by fitting with a single Gaussian function and the obtained fingerprint showed nine separated peaks, one for each donor-labeled nucleotide (**Figure 3 B-J**).

**Figure 3:**
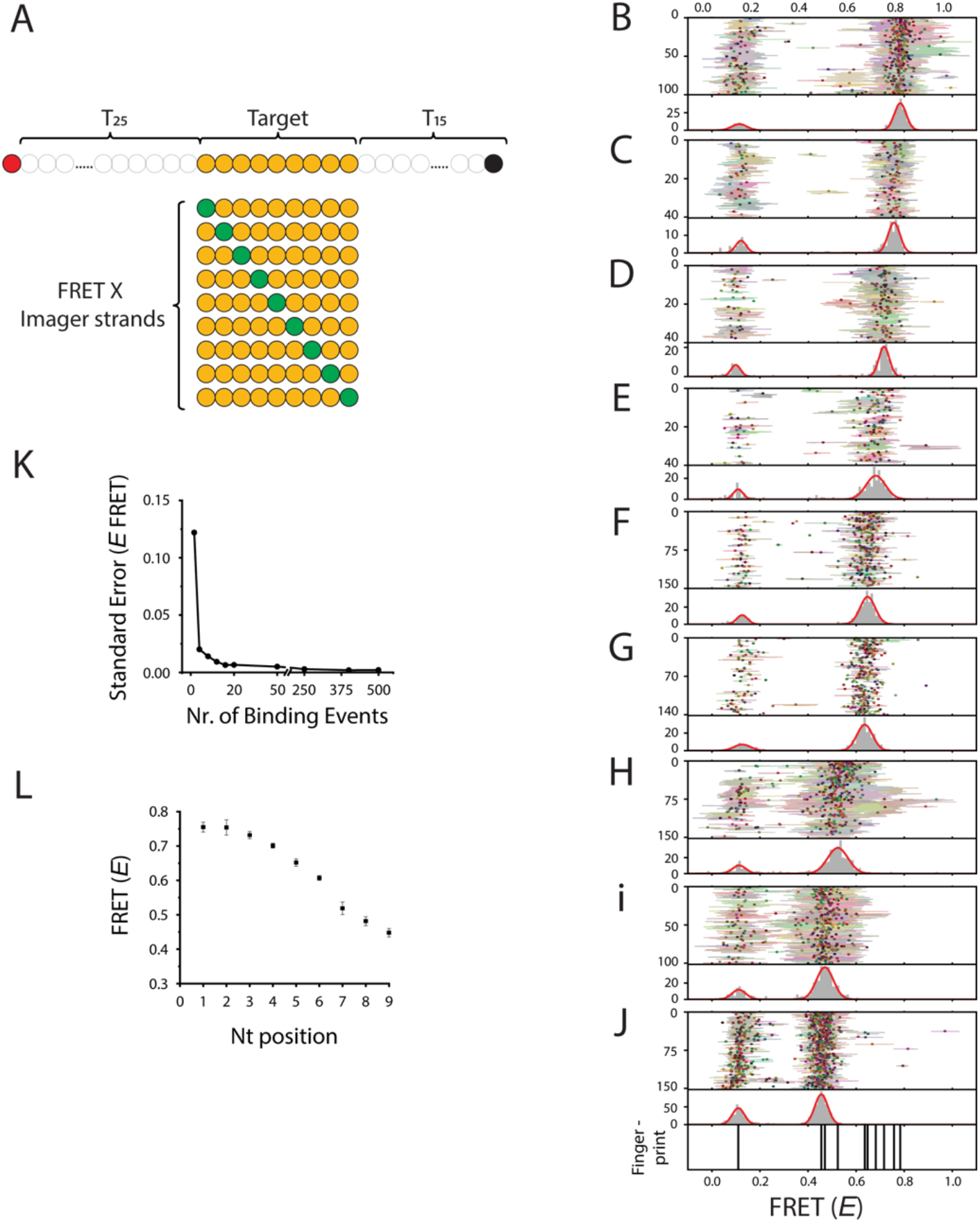
Single nucleotide resolution can be achieved with FRET X. A) Schematic representation of the single-molecule constructs used for the determination of different POIs separated by a single base pair. An acceptor (Cy5, red circle) labeled ssDNA target construct consisting of a 9 nt target sequence (orange circles) where each imager strand can bind. A series of donor (Cy3, green circles) labeled FRET X imager strands. The position of each POI (or nucleotide) in the target sequence will be determined one by one using our FRET X approach. B-J) Kymographs for each of the POIs determined using FRET X. The top kymograph is obtained with the imager strand where the donor fluorophore binds closest to the acceptor (separated by a 25-thymine linker). Each next kymograph is ontained with a subsequent imager strand where the distance to the acceptor increases by a single base pair. We obtained nine separated FRET histograms, one for each of donor labeled base pairs using FRET X. The bottom panel shows nine clearly separated peaks in the FRET fingerprint. The fingerprint shows the center of each Gaussian fit that was obtained using our FRET X approach. K) Standard error of the FRET X efficiency for imager strand 5 (Fig. 3F) vs the number of binding events. We observe that we can determine the center of a Gaussian fit with a FRET X precision of ΔE ~ 0.01 after >10 binding events. L) The mean FRET X efficiency for each of the POIs determined on different days. We find good reproducibility for FRET X. Mean FRET efficiencies and standard deviation are calculated from 3 independent experiments.

To determine the precision that can be obtained using our FRET X approach, the standard error of the FRET efficiency was plotted as a function of the number binding events. The chosen events were from an imager strand labeled at position 5 (**Figure 3F**) that yielded a FRET efficiency value of 0.65. We found that the center of a Gaussian fit can be determined with a precision of 1%p (or Δ*E*~0.01) after obtaining >10 binding events (**Figure 3K**). The reproducibility of FRET X was demonstrated by measuring all nine labeled imager strands on different days. As shown in **Figure 3L**, the standard deviation between the measurements made on different days is about 2%p per each construct.

Finally, to demonstrate the potential to use FRET X for structure analysis at the single molecule level, we designed two ssDNA constructs with structural differences and tested whether individual molecules can be distinguished when the two are randomly mixed. The ssDNA constructs consist of two POIs, one of which is located at an identical position on the two DNA constructs. The second POI is connected to the side of one of the nucleotides in the backbone sequence and has a different location on the two constructs (**Figure 4A and Supplementary Figures 4 and 5**). To avoid the photobleaching of the acceptor dye, we designed a unique sequence near the 3’ end of the construct where a complementary acceptor labeled imager strand can transiently bind. We immobilized a mixture of the two constructs with 1:1 ratio.

**Figure 4:**
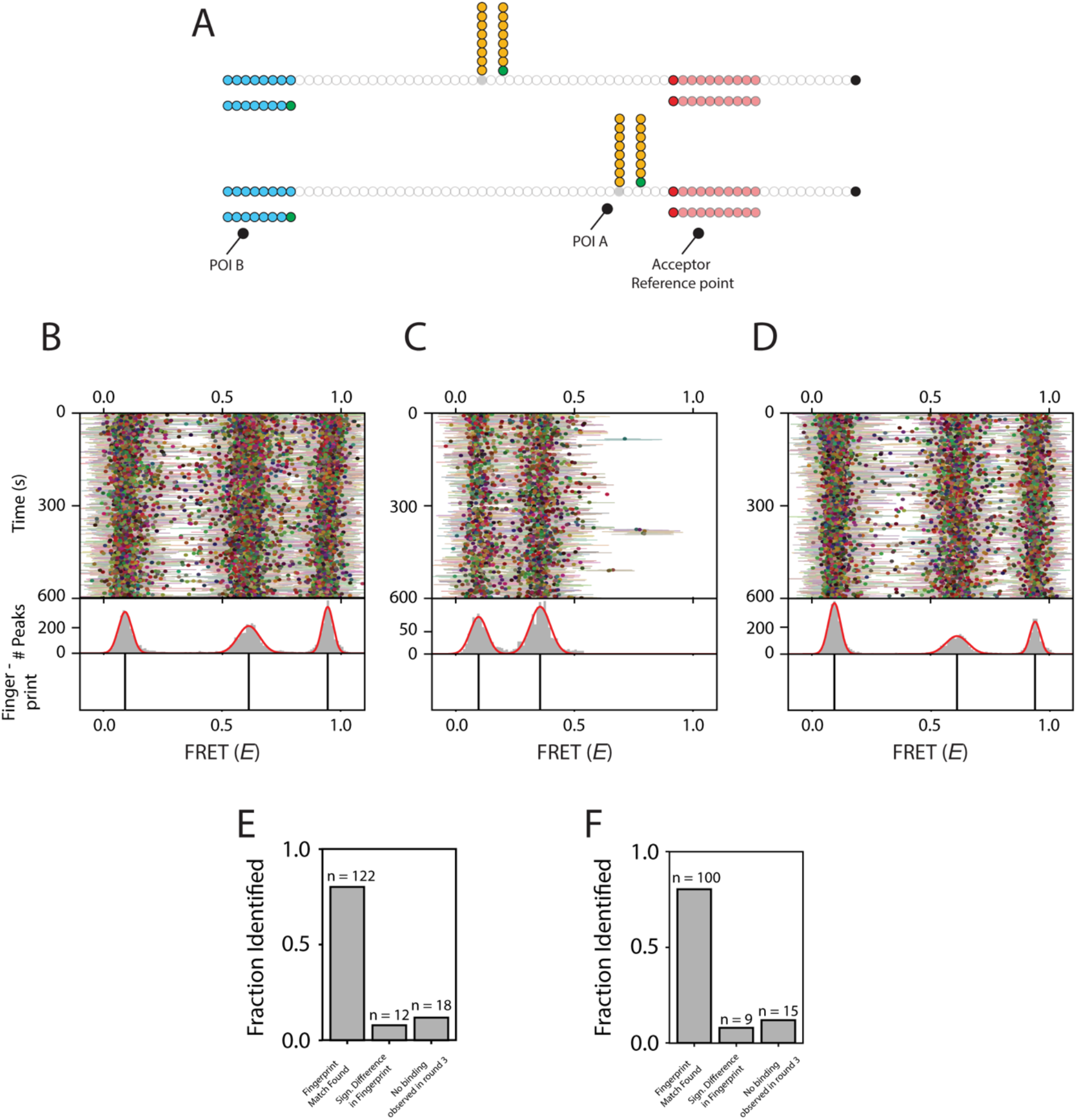
Structural analysis of individual molecules using FRET X. Schematic representation of the DNA constructs used for structural analysis. The ssDNA construct contains two POIs, of which one is fixed and has the same location relative to the acceptor on both constructs. The second POI is connected to the side chain of one of the nucleotides in the backbone sequence and has a different location on both constructs. B-D) Kymographs obtained for an equal mixture of the ssDNA constructs immobilized on the slide surface. The FRET X cycle consisted of 3 rounds. B) In a first round of the FRET X cycle, we observed two FRET peaks reporting on the distance of POI A from the acceptor for both constructs. C) The second round of the FRET X cycle resulted in a single peak obtained from FRET between POI B and the acceptor, which is identical in both constructs. D) The last round of the FRET X cycle we confirmed the location of POI A and observed the same FRET peaks as in round1. E and F) Bar plots showing the fractions of fingerprint matches and non-matches for individual molecules that were identified as medium-(Fig. 4E) or high-FRET construct (Fig. 4F). We determined the mean FRET efficiency of the medium- or high-FRET fingerprint in round 1 and compared this with a detection uncertainty of Δ*E* ~ 0.07 with round 3 to find positives matches. The majority of molecules were identified identically between round 1 and 3, for the medium-(Fig. 4E) and high-(Fig. 4F) FRET ssDNA constructs.

In a first round of FRET X, we determined the FRET efficiency for POI A and observed two distinct FRET populations reporting on the distinct distance between POI A and the acceptor reference point, depending on the constructs (**Figure 4B**). Next, we washed the microfluidic chamber and injected the imager strand for POI B. As expected, we observed a single peak for POI B, reporting on the similar position of POI B for both constructs (**Figure 4C**). In a final round of FRET X, we confirmed the location of POI A by agin injecting the imager strand for POI A back and observed the same FRET peaks as in the FRET X imaging rounds 1 (**Figure 4 B and D**).

For each individual molecule, we determined the mean FRET efficiency for POI A in round 1 and compared this with the FRET efficiency obtained for POI A in round 3 (**Supplementary Figure 6**). The majority (>80 %) of the individual molecules in the mixture had a similar resolved FRET efficiency of POI A between rounds 1 and 3, for the medium-(**Figure 4E**) or high-(**Figure 4F**) FRET constructs. Only a small fraction of molecules did not show a match between the FRET X rounds due to a different resolved FRET efficiency for POI A or a lack of imager strand binding events (**Figure 4E and F**). Altogether, these results show that the FRET X method is capable of detecting the structure of individual DNA constructs at the single-molecule level.

## DISCUSSION

Here we present a proof-of-concept for FRET X, a novel tool for the detection of several FRET pairs in a single object, which can be used for the structural analysis of biomolecules. Our FRET X technique relies on the dynamic binding of short fluorescently labeled oligos to complementary docking sequences on a target object. Conventional single-molecule FRET techniques report on the changes in distance between a single dye pair on a single molecule. Multicolor FRET approaches have been developed, but they are often difficult to implement due to the complex sample preparation and complicated analysis of multispectral fluorescence signals. In contrast, FRET X uses orthogonal imager strands for different POIs allowing for the detection of a large number of POIs on a single object.

Conventional single-molecule FRET analysis of protein structures is labour intensive as protein molecules need to be modified for site-selective labeling. We demonstrated that FRET X can be a tool for the structural analysis of complex biomolecules, since it can report on the distances of multiple POIs on a single biomolecule. By covalently attaching short ssDNA strands, which act as FRET X docking strands, to various regions of interest on a protein, one will be able to detect a large number of residues or domains in a single experiment. For example, for the analysis of a protein structure, the attachment of FRET X docking strands can be attached by using orthogonal chemistry for surface-exposed cysteine or lysine residues in proteins. This method eliminates the need for any complicated preparative engineering of the biomolecules for site-selective labeling.

## MATERIALS AND METHODS

### Single-Molecule Setup

All experiments were performed on a custom-built microscope setup. An inverted microscope (IX73, Olympus) with prism-based total internal reflection was used. In combination with a 532 nm diode-pumped solid-state laser (Compass 215M/50mW, Coherent). A 60x water immersion objective (UPLSAPO60XW, Olympus) was used for the collection of photons from the Cy3 and Cy5 dyes on the surface, after which a 532 nm long pass filter (LDP01-532RU-25, Semrock) blocks the excitation light. A dichroic mirror (635 dcxr, Chroma) separates the fluorescence signal which is then projected onto an EM-CCD camera (iXon Ultra, DU-897U-CS0-#BV, Andor Technology). A series of EM-CDD images was recorded using custom-made program in Visual C++ (Microsoft).

### Single-Molecule Data Acquisition

Single-molecule flow cells were prepared as previously described.^20,21^ In brief, to avoid non-specific binding, quartz slides (G. Finkerbeiner Inc) were acidic piranha etched and passivated twice with polyethylene glycol (PEG). The first round PEGylation was performed with mPEG-SVA (Laysan Bio) and PEG-biotin (Laysan Bio), followed by a second round of PEGylation with MS(PEG)4 (ThermoFisher). After assembly of a microfluidic chamber, the slides were incubated with 20 μL of 0.1 mg/mL streptavidin (Thermofisher) for 2 minutes. Excess streptavidin was removed with 100 μL T50 (50mM Tris-HCl, pH 8.0, 50 mM NaCl). Next, 50 μL of 75 pM Cy5 labeled ssDNA was added to the microfluidic chamber. After 2 minutes of incubation, unbound ssDNA was washed away with 100 μL T50. For experiments in Figure 1, 50 μL of 10 nM donor labeled imager strands in imaging buffer (50 mM Tris-HCl, pH 8.0, 500 mM NaCl, 100 mM MgCl_2_, 0.8 % glucose, 0.5 mg/mL glucose oxidase (Sigma), 85 ug/mL catalase (Merck) and 1 mM Trolox (Sigma)) was injected. All single-molecule FRET experiments were performed at room temperature (23 ± 2 °C).

#### FRET X Imaging (Figures 2 and 3)

For FRET X imaging in Figures 2 and 3, 50 μL of 75 pM target DNA strands were immobilized and the unbound DNA was washed away with 100 μL T50 after 2 minutes of incubation. Next, an imaging buffer containing the imager strand for POI A (**Figure 2**), or imager strand with internal nucleotides labeled at position 1 (**Figure 3**), was injected. After obtaining 2000 frames at 100 ms exposure time, the microfluidic chamber was washed with 1000 μL T50 and the imager strand for POI B (**Figure 2**), or internally labeled nucleotide position 2 (**Figure 3**), was injected. This cycle was repeated until all internally labeled nucleotides were measured for Figure 3.

#### FRET X experiments for Single molecule Structural Analysis (Figure 4)

For buffer exchange and imaging of the same molecules in a single field of view for different rounds of FRET X imaging, tubing was connected to the inlet and outlet of the microfluidic chamber. One of the tubes was connected to a buffer reservoir and the other was connected to a syringe. By gently pulling on the syringe, the washing buffers and imaging solutions were exchanged without perturbing the sample stage.

For the branched DNA constructs experiments, 50 μL of 75 pM branched DNA target strand was immobilized for 2 minutes and unbound DNA was removed with 100 μL T50. To increase the probability of energy transfer between donor and acceptor fluorophores, the acceptor imager strand was designed to have a dissociation rate of ~ 0.1 s^-1^.^22^ For long term acquisition, a 50 μL imaging solution consisting of 100 nM acceptor imager strand and 10 nM of donor labeled imager strand for POI A was injected and the chamber was imaged for 15 minutes at 100 ms exposure time. Then the imaging solution for POI A was removed by washing with 1000 μL T50 and the imaging solution of POI B was added (50 μL of 100 nM acceptor imager strand and 10 nM of imager strand for POI B in imaging buffer). After this second round of imaging, the microfluidic chamber was washed with 1000 μL T50 and POI A was imaged again by injecting fresh imaging solution for POI A.

### Data Analysis

CCD images were analyzed using a custom code written in IDL (ITT Visual Information Solution) to find the position of individual FRET pairs and to extract fluorescence time traces. When the same field of view is measured multiple times (**Figure 4**), drift correction between the measurements and trace extraction were performed by a custom-built code written in Python (Python 3.7). For visualization of single molecule fluorescence and FRET time traces, we used a custom code written in Matlab (Mathworks). For automated detection of individual fluorescence barcode binding events, we used a custom Python code (Python 3.7) utilizing a two-state K-means clustering algorithm on the sum of the donor and acceptor fluorescence intensities of individual molecules to identify the frames with high intensities. ^[Kim et, al, in preperation]^ To avoid false positive detections, only binding events that lasted for more than three consecutive frames were selected for further analysis. FRET efficiencies for each barcode binding events were calculated and used to build the FRET kymograph and histogram. Populations in the FRET histogram are automatically classified by using Gaussian mixture modeling and used to determine the presence of specific barcodes of interest. The automated analysis code in Python is freely available at (https://github.com/kahutia/transient_FRET_analyzer2).

## ACKNOWLEDGEMENTS

We thank Viktorija Globyte for critical reading and feedback. C.J. was supported by Vrije Programma (SMPS) of the Foundation for Fundamental Research on Matter and Human Frontier Science Program (RGP0026/2019).

## AUTHOR CONTRIBUTIONS

M.F. and C.J. initiated and designed the project. M.F. and S.H.K. performed the experiments. S.H.K and I.S. wrote the analysis software. M.F., S.H.K., C.J. analysed and discussed the data. M.F. and C.J. wrote the manuscript. All authors read and improved the manuscript.

