## Supplementary Information for "High-Resolution Single-Molecule FRET via DNA eXchange (FRET X)"

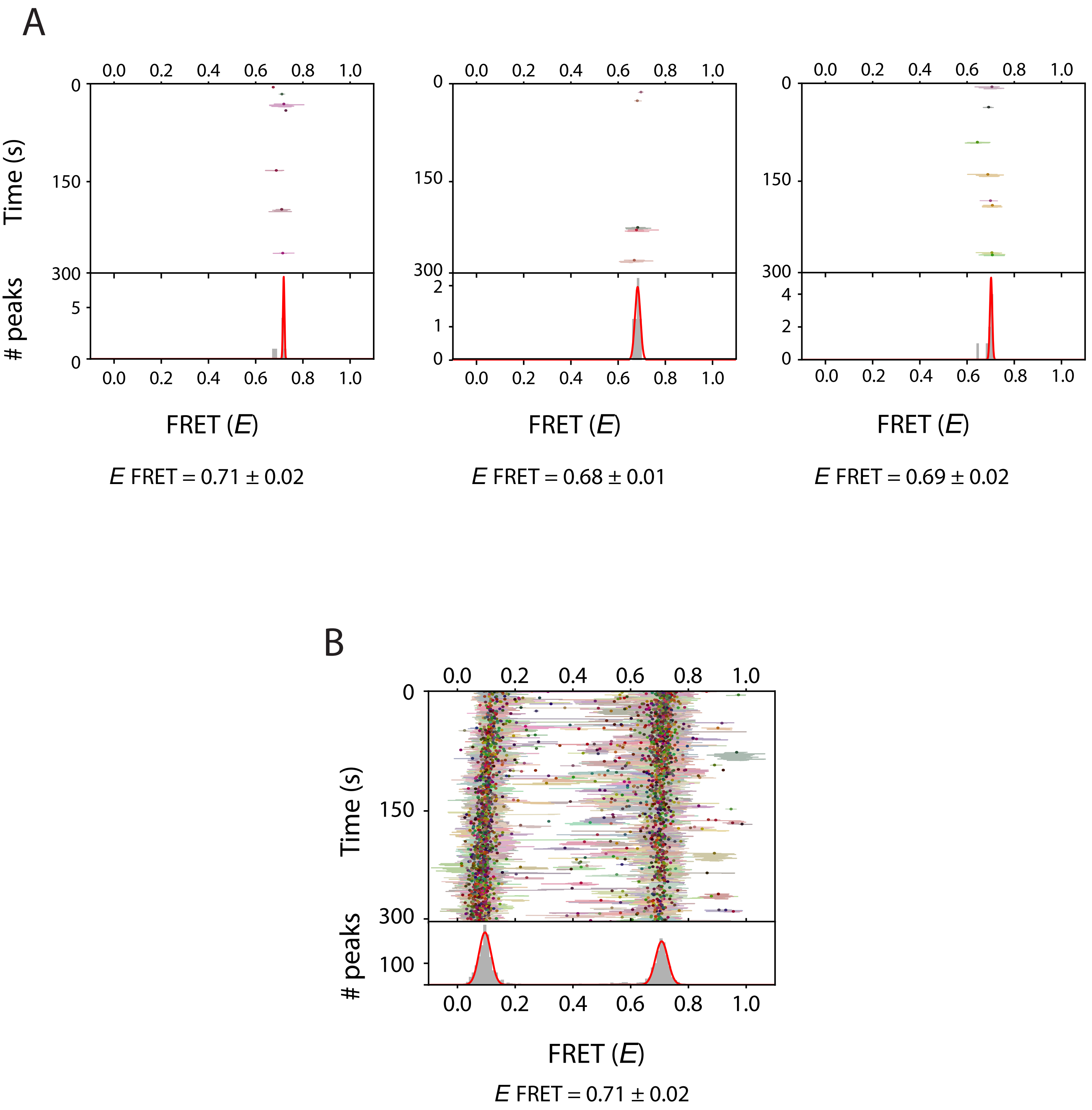


Supplementary Figure 1: Single-Molecule FRET Kymographs.
A) Representative FRET kymographs obtained from individual molececule in a single field of view.
B) An ensemble FRET kymograph obtained from all molecules in a single field of view.
The FRET efficiencies were reported as the mean ± the standard deviation.

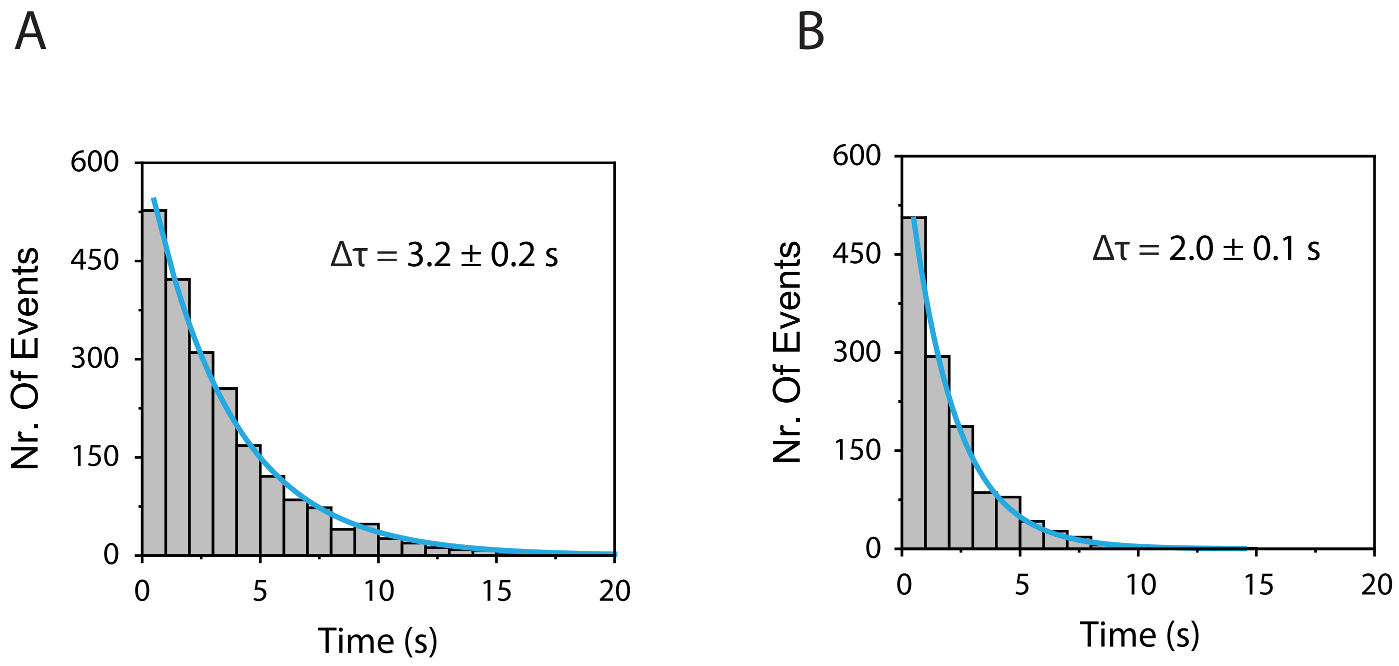


Supplementary Figure 2: Single-molecule binding kinetics of FRET X imager strands
A) Dwell-time histogram for FRET X imager strands used for detection of POI A in Figure 2. Maximum likelood estimation gives 3.2 ± 0.2 s for a single exponential distribution (blue line). The number of datapoints:  n = 2146.
B) Dwell-time histogram for FRET X imager strands used for detection of POI B in Figure 2. Maximum likelood estimation gives 2.0 ± 0.1 s for a single exponential distribution (blue line). The number of datapoints: n = 1252.

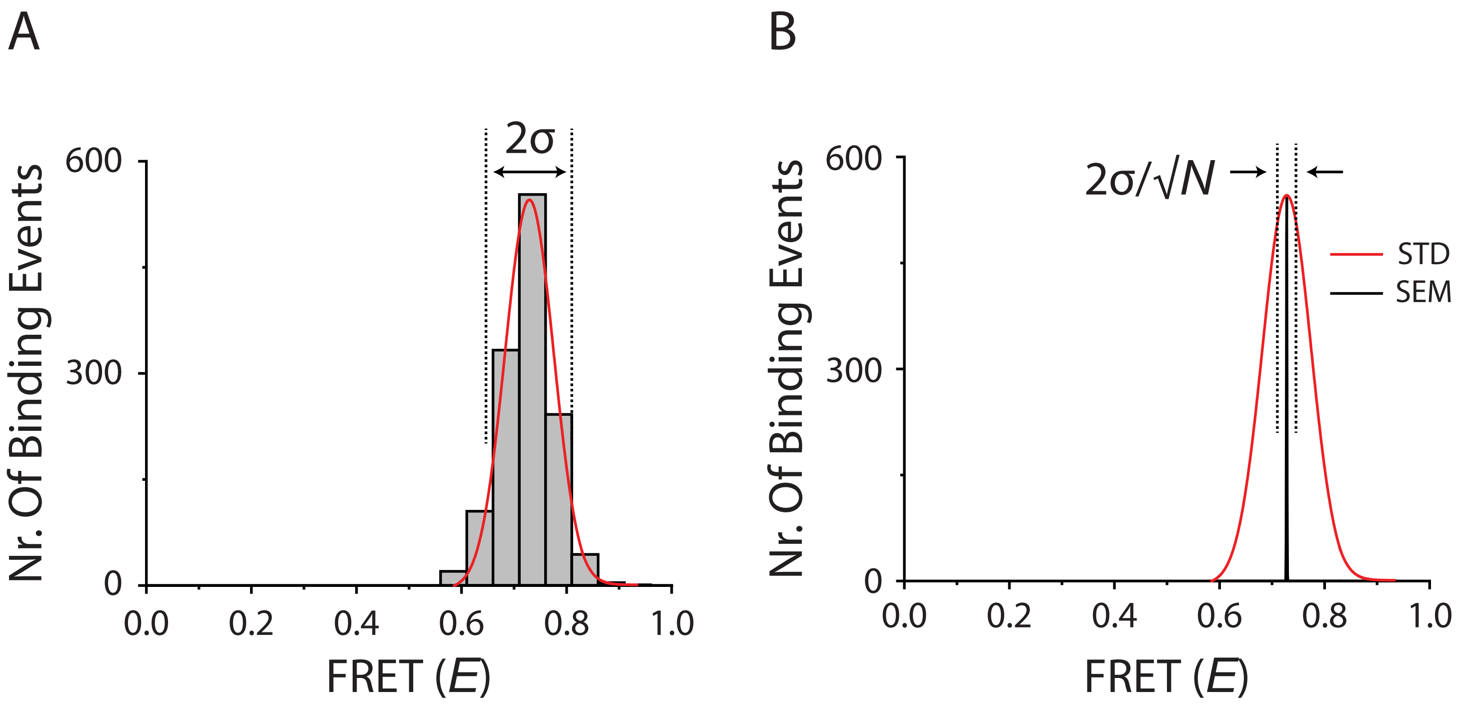


Supplementary Figure 3: Standard deviation vs Standard error
A) The standard deviation (σ) reports on the intrinstic broadness of a FRET histogram.
B) The standard error measures the accuracy of determing the center of a peak. N is the number of imager strand binding events.

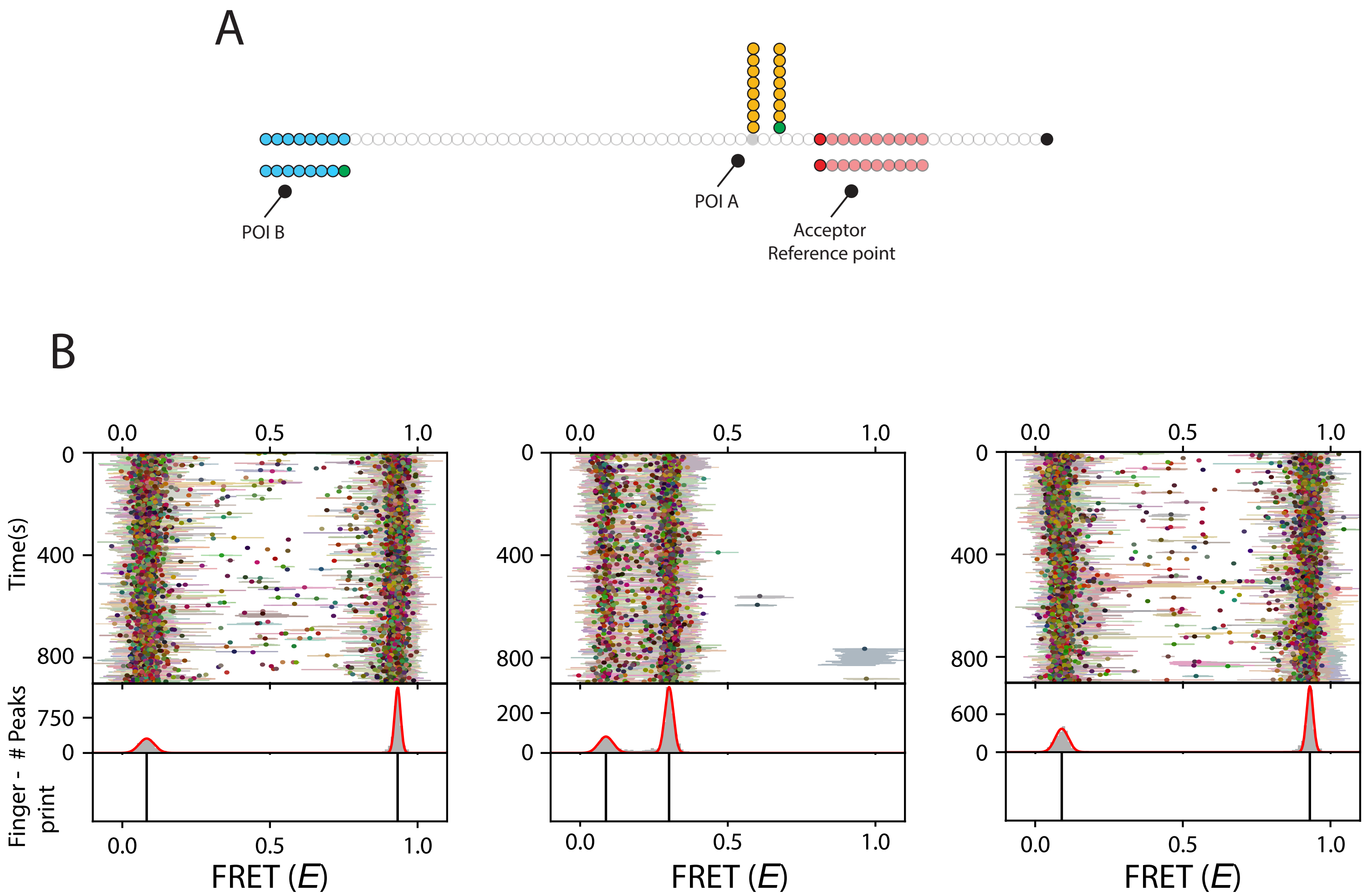


Supplementary Figure 4: Single-Molecule FRET X analysis of complex ssDNA structure resulting in high FRET.

A) Schematic representation of complex ssDNA structure resulting in high FRET. Upon binding of the FRET X imager strand for POI A, the donor fluorophore is separated by a 5 nt polyT linker from the acceptor binding site. The imager strand for POI B is separated by a 40 nt polyT linker from the acceptor binding site.

B) Ensemble FRET kymographs obtained from different rounds of FRET X imaging. In a first round of imaging (left panel) we obtained a high FRET peak reporting on the location of POI A relative to the acceptor biding site. After washing of the microfluidic cell we injected the imager strand for POI B (middle panel) and obtained a low FRET efficiency reporting on the distance of POI B to the acceptor binding site. In a last round of FRET X imaging (right panel) we confirmed the high FRET peak for POI A relative to the acceptor binding site.

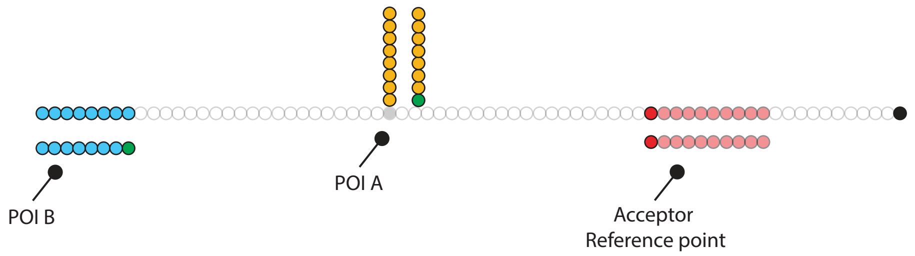


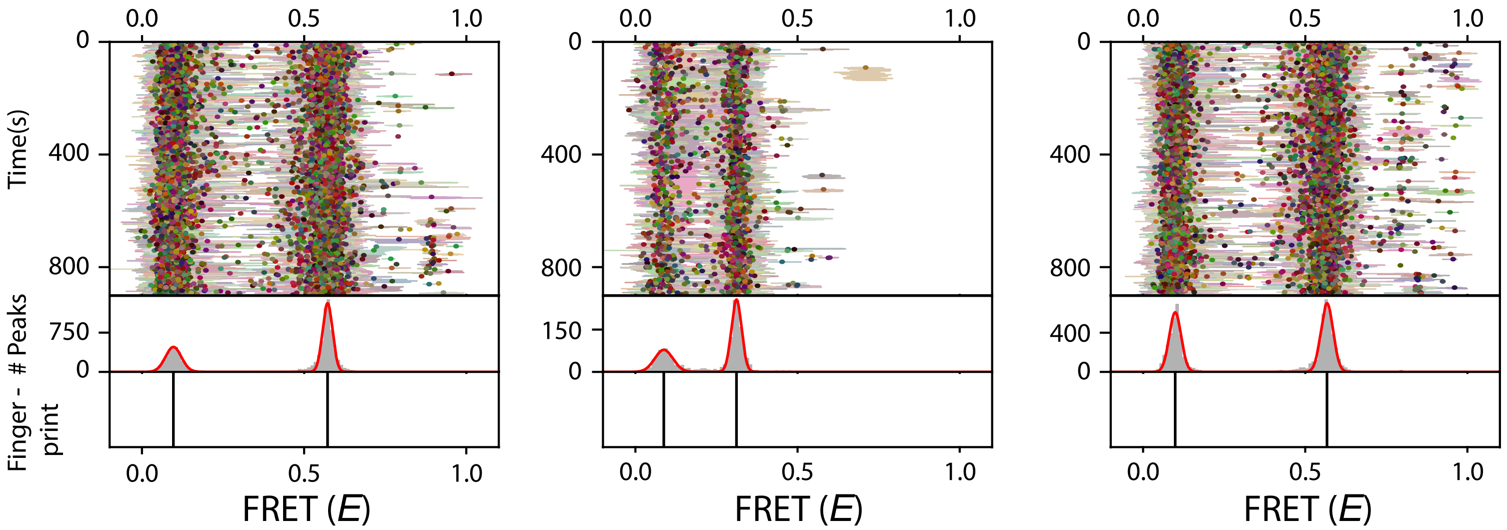


Supplementary Figure 5: Single-Molecule FRET X analysis of complex ssDNA structure resulting in medium FRET

A) Schematic representation of complex ssDNA structure resulting in medium FRET. Upon binding of the FRET X imager strand for POI A, the donor fluorophore is separated by a 20 nt polyT linker from the acceptor binding site. The imager strand for POI B is separated by a 40 nt polyT linker from the acceptor binding site.

B) Ensemble FRET kymographs obtained from different rounds of FRET X imaging. In a first round of imaging (left panel) we obtained a medium FRET peak reporting on the location of POI A relative to the acceptor biding site. After washing of the microfluidic cell we injected the imager strand for POI B (middle panel) and obtained a low FRET efficiency reporting on the distance of POI B to the acceptor binding site. In a last round of FRET X imaging (right panel) we confirmed the medium FRET peak for POI A relative to the acceptor binding site.

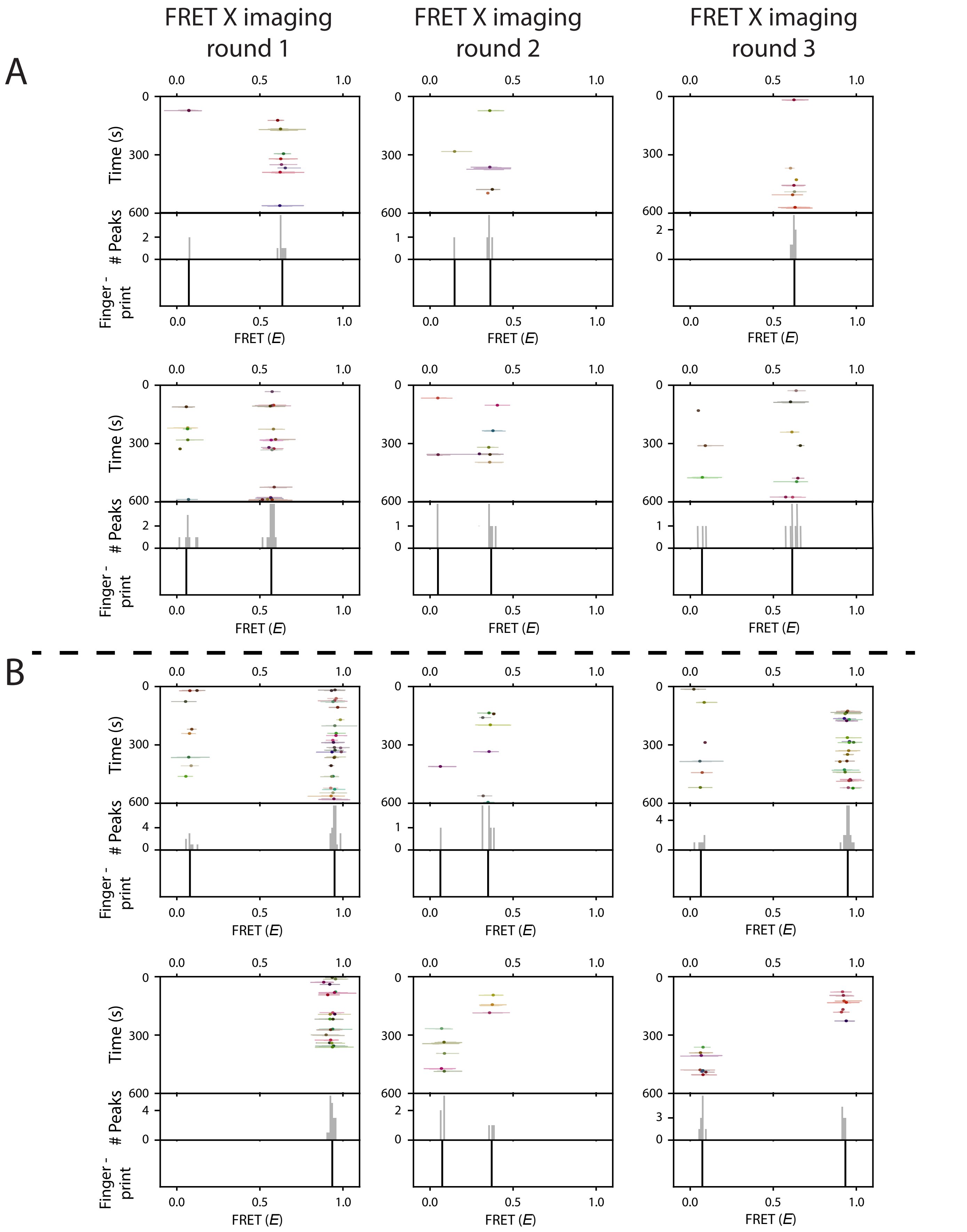


Supplementary Figure 6: Representative kymographs of individual molecules in a mixture of structurally similar DNA constructs
A-B) Representative FRET kymographs from individual complex ssDNA molecules in a mixture. Using our FRET X approach we can observe a difference between the medium FRET complex structure (Fig. S-6A) or high FRET complex structure (Fig. S-6B), in FRET X imaging round 1 (left Column) and round 3 (right column). The second round of FRET X imaging (middle column) shows a similar FRET peak for both constructs, reporting on the structural similarity among the constructs.
